# Kilosort: realtime spike-sorting for extracellular electrophysiology with hundreds of channels

**DOI:** 10.1101/061481

**Authors:** Marius Pachitariu, Nicholas Steinmetz, Shabnam Kadir, Matteo Carandini, Harris Kenneth D.

## Abstract

Advances in silicon probe technology mean that in vivo electrophysiological recordings from hundreds of channels will soon become commonplace. To interpret these recordings we need fast, scalable and accurate methods for spike sorting, whose output requires minimal time for manual curation. Here we introduce Kilosort, a spike sorting framework that meets these criteria, and show that it allows rapid and accurate sorting of large-scale in vivo data. Kilosort models the recorded voltage as a sum of template waveforms triggered on the spike times, allowing overlapping spikes to be identified and resolved. Rapid processing is achieved thanks to a novel low-dimensional approximation for the spatiotemporal distribution of each template, and to batch-based optimization on GPUs. A novel post-clustering merging step based on the continuity of the templates substantially reduces the requirement for subsequent manual curation operations. We compare Kilosort to an established algorithm on data obtained from 384-channel electrodes, and show superior performance, at much reduced processing times. Data from 384-channel electrode arrays can be processed in approximately realtime. Kilosort is an important step towards fully automated spike sorting of multichannel electrode recordings, and is freely available (github.com/cortex-lab/Kilosort).

## 1 Introduction

The oldest and most reliable method for recording neural activity involves lowering an electrode into the brain and recording the local electrical activity around the electrode tip. Action potentials of single neurons generate a stereotypical temporal deflection of the voltage, known as a spike waveform. When multiple neurons close to the electrode fire action potentials, their spikes must be identified and assigned to the correct neuron based on the features of the recorded waveforms, a process known as spike sorting^1–15^.

Measuring voltage at multiple closely-space sites in the extracellular medium can substantially improve spike sorting accuracy. In this case, the recorded waveforms also have characteristic spatial shapes, determined by each neuron’s location and physiological characteristics. Together, the spatial and temporal shape of the waveform provide all the information that can be used to assign a given spike to a neuron^16^.

Current methods for spike sorting, however, will struggle to meet the requirements raised by a new generation of high-count, high-density electrodes that are soon to become commonplace. These electrodes have several hundred closely-spaced recording sites^17^, and initial tests suggest that they can reveal the activity of 100 to 1,000 neurons firing tens of millions of spikes. When applied to such data, algorithms designed for tens of recording sites^18,19^ suffer from substantial limitations. The automatic sorting software can take days to weeks to run, and require hours to days of manual curation. Furthermore, with more channels and higher density, resolution of spatiotemporally overlapping spikes becomes both more tractable and more important.

**Figure 1.**
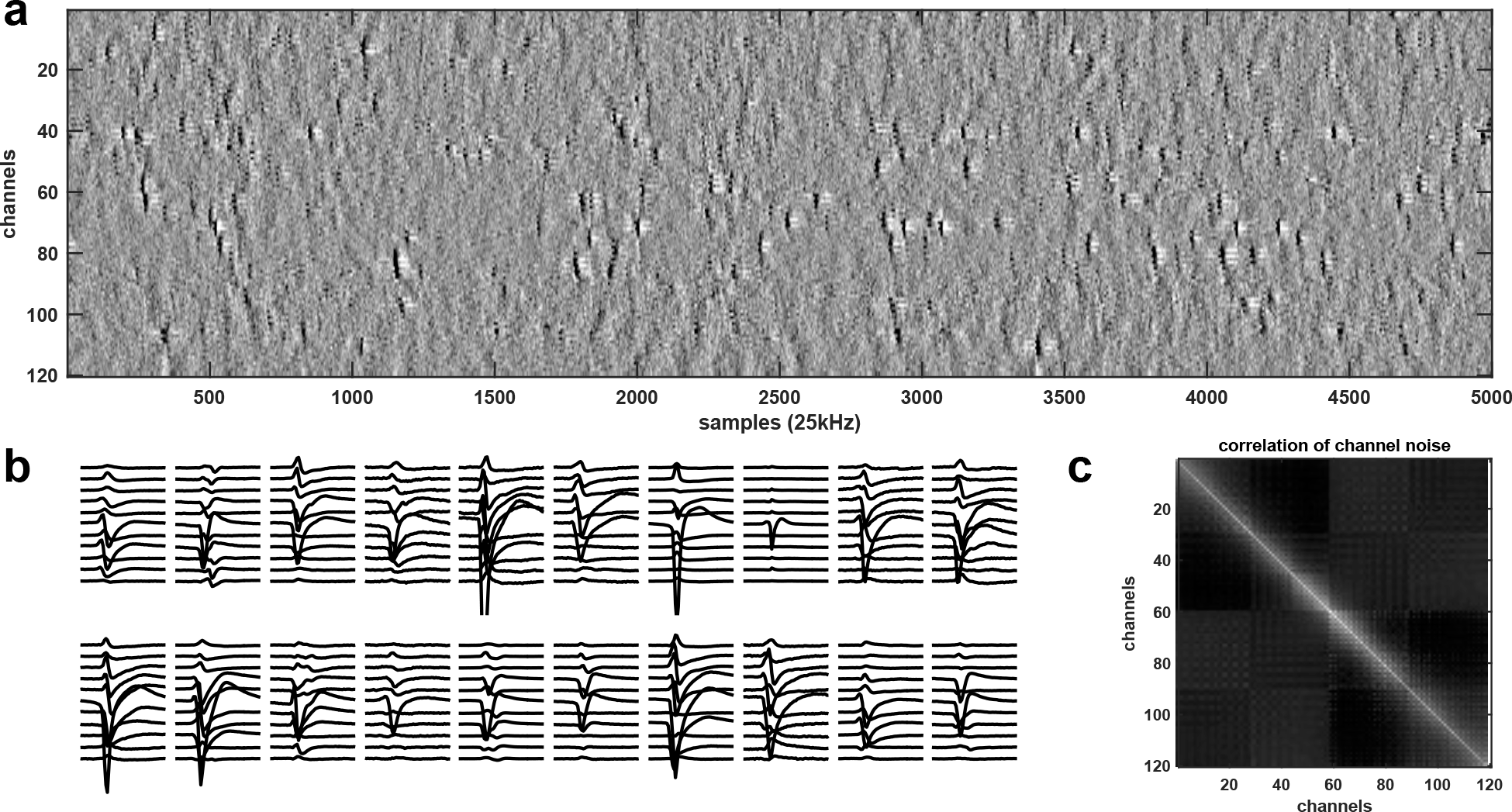
Data from high-channel count recordings. **a,** High-pass filtered and channel-whitened data. Negative peaks are action potentials. **b,** Example mean waveforms, centered on their peaks. **c,** Example cross-correlation matrix across channels (before whitening, no spikes included).

Here we overcome these limitations and present Kilosort, a new algorithm which takes advantage of a novel mathematical approach that greatly reduces the amount of calculation required, together with the computing capabilities of low-cost commercially available graphics processing units (GPUs). To illustrate its abilities we show that it accurately spike sorts the output of 384-channel dense probes in approximately real time.

### *1.1* *High-density electrophysiology and structured sources of noise*

With high-density neural probes (i.e. site spacing in the range ~20 *μ*m), the waveforms of each neuron can be typically detected on 5 to 50 channels simultaneously (Fig. 1a,b; example data available at http://data.cortexlab.net/dualPhase3). This provides a substantial amount of information per spike, but because other neurons also fire on the same channels, a clustering algorithm is required to unmix the signals and assign spikes to the correct neuron. Furthermore, structured sources of noise can make this assignment more difficult. For example, neurons that are too distant from the electrode to be sortable provide myriad superimposed spike waveforms, a continuous random background against which the features of sortable spikes must be distinguished^16^.

### *1.2* *Previous work*

A traditional approach to spike sorting divides the problem into several stages^2^. First, spike times are detected, for example as times when the negative voltage crosses a pre-defined threshold. Second, these spike waveforms are extracted and projected into a common low-dimensional space, typically obtained by principal component analysis (PCA^20^. Third, the spikes are clustered in this low-dimensional space using a variety of approaches, such as mixtures of Gaussians^18^ or peak-density detection^21^. Some algorithms also include a fourth stage of template matching that scans the raw data for overlapping spikes, which may have been missed in the first detection phase^11,12,14^. Finally, a manual curation stage is required, in which a human operator corrects the imperfect automated results using a graphical user interface (GUI). This last step is particularly necessary for recordings subject to electrode drift, where the waveforms of a given neuron vary over time and may be assigned to multiple clusters.

Here we describe a system that omits spike detection and PCA and instead combines the identification of template waveforms and associated spike times in a single unified model. This model seeks to reconstruct the entire raw voltage dataset with the templates of candidate neurons. We define a cost function for this reconstruction, and derive approximate inference and learning algorithms that can be successfully applied to very large channel count data. This approach is related to a previous method^6^, but that method requires a generic convex optimization that is slow for recordings with large numbers of channels.

As we demonstrate with constructed ground-truth datasets, our system is more accurate than a current widely-used method^18^. Furthermore, we demonstrate that on real datasets with 384 channels, this implementation is fast enough to run in nearly real time.

## 2 Model formulation

We start with a generative model of the raw electrical voltage. Unlike the traditional pipeline, this algorithm does not start with a spike detection step, nor project the spike waveforms to a lower-dimensional PCA space. As we show below, both of these steps would discard potentially useful information.

### *2.1* *Pre-processing: common average referencing, temporal filtering and spatial whitening*

To remove low-frequency fluctuations, such as the local field potential, we high-pass filter each channel of the raw data at 300 Hz. To diminish the effect of artifacts shared across all channels, we subtract at each timepoint the median of the signal across all recording sites, an operation known as common average referencing^22^. This step is best performed after high-pass filtering, because the LFP magnitude is variable across channels and can be comparable in size to the artifacts.

Next, we whiten the data across channels to remove correlated noise. In the frequency range typical of spikes, spatially correlated noise arises primarily from neurons far from the probe, whose spikes are too small to sort directly^16,23^ and have a large spatial spread over the surface of the probe. Since there are many such neurons, their noise averages out to have a stereotypical crosscorrelation pattern across channels (Fig. 1c). To estimate this noise covariance, we first remove the times of putative spikes (detected with a threshold criterion). We then estimate the covariance matrix ∑, and use its singular vectors and singular values *E, D* to obtain a symmetrical whitening matrix that maintains the spatial structure of the data, known as zero-phase component analysis (ZCA): *W*_ZCA_ = ∑^−1/2^ = *ED*^−1/2^*E*^*T*^. To regularize *D*, we add a small value to its diagonal. For very large channel counts, estimation of the full covariance matrix ∑ is noisy, and we therefore compute the columns of the whitening matrix *W*_ZCA_ independently for each channel, based on its nearest 32 neighbors. We then multiply the raw data matrix containing all channels with this whitening matrix.

**Figure 2.**
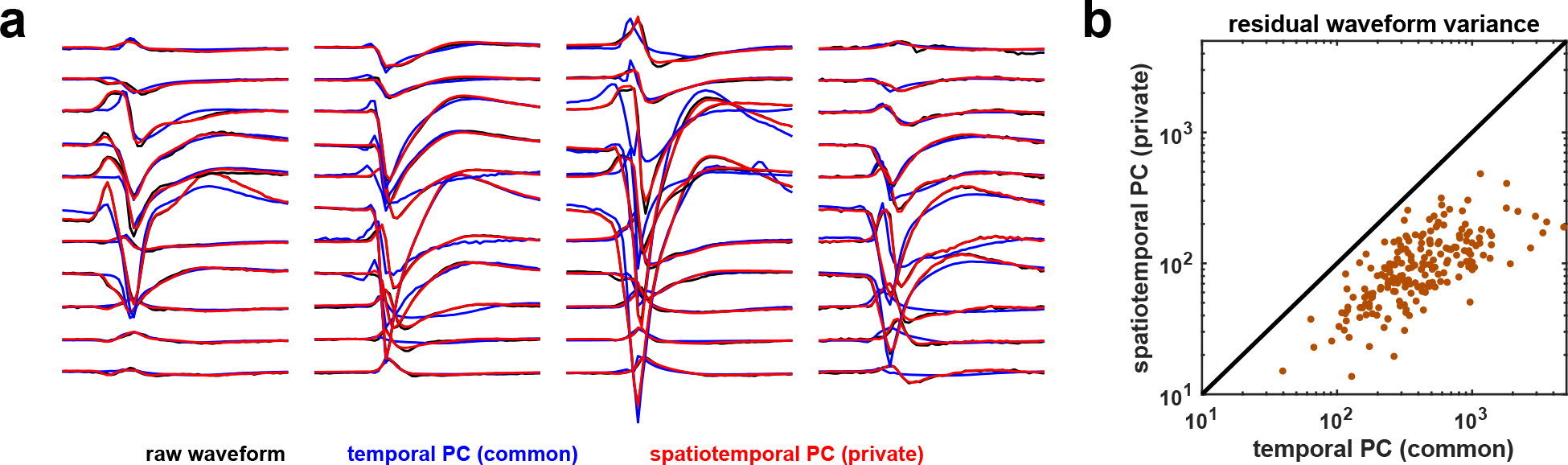
Spike reconstruction from three private PCs. **a,** Four example average waveforms (black) with their respective reconstruction from three common temporal PCs/channel (blue), and with reconstruction from three spatiotemporal PCs private to each spike (red). The red traces mostly overlap the black traces. **b,** Summary of residual waveform variance for all neurons in one dataset.

### *2.2* *Modelling mean spike waveforms with SVD*

When single spike waveforms are recorded across a large number of channels, most channels will have no signal and only noise. To prevent the large total energy on these many noise channels from swamping the signal present on a the smaller number of signal channels, previous approaches have estimated a “mask” to exclude channels with insufficient SNR to identify any given spike^18,19^; to further reduce noise and lower dimensionality, the spikes are usually projected into a small number of temporal principal components per channel^20^, typically three.

Here we introduce a different method for simultaneous spatial denoising/masking and for lowering the dimensionality of spikes. This method is based on the observation that any mean spike waveforms can be well explained by a singular value decomposition (SVD) decomposition of its spatiotemporal waveform, with as few as three components, but that the spatial and temporal components required can vary substantially between neurons (Fig. 2a). This approach of tailoring “private PCs” to each spike allows us to fit the spikes with ~5 times less residual variance than the standard approach of applying a single PCA approximation per channel, to all neurons on that channel (Fig. 2b). This decomposition results in an automated masking strategy, which allows the waveforms to be denoised and irrelevant channels ignored, and also speeds up the algorithm, by allowing the use of standard low-rank filtering techniques (see below).

### *2.3* *Integrated template matching framework*

To define a generative model of the electrical recorded voltage, we take advantage of the approximately linear summation of electrical potentials from different sources in the extracellular medium. We combine the spike times of all neurons into a *N*_spikes_-dimensional vector s, such that the waveforms start at time samples s + 1. We define the cluster identity of spike *k* as σ(*k*), taking values into the set {1,2,3,…,*N*}, where *N* is the total number of neurons. We represent the normalized waveform of neuron *n* as the matrix *K_n_* of size number of channels by number of sample timepoints *t_s_* (typically 61). The matrix *K_n_* is approximated by a three-dimensional singular value decomposition, *K_n_* = *U_n_W_n_*, whereby *K_n_* is deconstructed into three pairs of spatial and temporal basis functions, *U_n_* and *W_n_*, such that the norm of *U_n_W_n_* is 1. The value of the electrical voltage at time *t* on channel *i* is modeled by

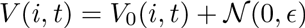

where the noise is modelled as independent Gaussian of variance *ϵ*.*V*_0_(*I*,*t*) is defined as

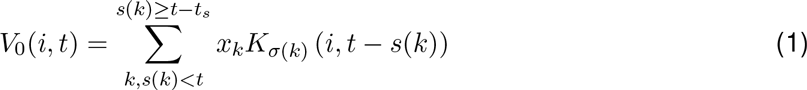

where the index *k* picks out those spikes that overlap with the timepoint *t*, because they happen at nearby times *s*(*k*), and *x_k_* > 0 is the amplitude of spike *k*, further constrained by

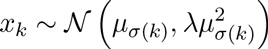

This last equation models variations in spike amplitudes for spikes from the same neuron, due to factors like burst adaptation and drift. We modelled this variability with a Gaussian distribution whose variance scales with the square of its mean, to capture the fact that the spikes of neurons closer to the probe vary in relative, not absolute amplitude. λ and *ϵ* are hyperparameters that control the relative scaling with respect to each other of the reconstruction error and the prior on the amplitude.

This model formulation leads to the following cost function, which we minimize with respect to spike times *s*, spike amplitudes *x*, templates *K*, and cluster assignments *σ*:

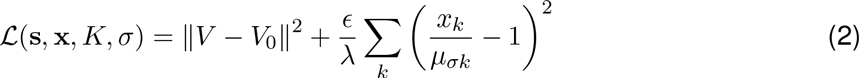

The second term in this expression has the purpose of limiting the number of spikes that are assigned amplitudes that deviate strongly from the mean of the relevant cluster. It is scaled by the ratio 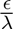, which we usually set to a constant between 1 and 10.

## 3 Learning and inference in the model

To optimize the cost function, we first initialize the templates and then alternate between two steps: finding the best spike times s, cluster assignments *σ*, and amplitudes x (template matching); and optimizing the template waveforms *K* for a given s, *σ*,x (template optimization). After the final spike times and amplitudes have been extracted, we run a final post-optimization merging algorithm which finds pairs of clusters whose spikes form a single continuous density. These steps are described in detail below.

### *3.1* *Stacked initializations with prototypical spikes and scaled K-means*

The density of spikes can vary substantially across the probe, depending on the location of each recording site in the brain. It is thus helpful to initialize the algorithm in a way that matches the number of clusters to local spike density. This approach reduces the need for moving templates from one part of the probe to another during the optimization process, which would be prone to local minima. To initialize the templates, we thus start by detecting spikes using a threshold rule, and progress through the recording keeping a running subset of prototypical spikes that are sufficiently different from each other by an L2 norm criterion. During this initialization phase, we excluded overlapping spikes by enforcing a minimum spatiotemporal peak isolation criterion. Out of the prototypical spikes thus detected, we select the top *N* (number of desired clusters) which had most matches to other spikes in the recording.

These *N* prototypes are then used to initialize a scaled K-means algorithm. This algorithm uses the same cost function described in equation 2, with spike times s fixed to those found by a threshold criterion. Note that unlike standard K-means, each spike is allowed to have variable amplitude^24^, which allows initialization to proceed robustly even though amplitudes are variable.

### *3.2* *Inferring spike times and amplitudes via template matching*

The inference step attempts to find the best spike times, cluster assignments and amplitudes, given a set of templates {*K_n_*} with low rank-decompositions *K_n_* = *U_n_W_n_* and mean amplitudes *μ_n_*. After processing every hundred batches (or more, depending on their time length), the templates are obtained from the running average waveform *A_n_*, using an SVD decomposition to give *A_n_* ~ *μ_n_K_n_* = *μ_n_U_n_W_n_*, with 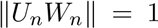, where *U_n_* is orthonormal and *W_n_* is orthogonal but not normalized. The primary roles of the low-rank representation are to guarantee fast inferences and to regularize and mask the waveform model.

Spike time inference during the iterative optimization stage is achieved with a standard template matching algorithm. The algorithm finds times *t* at which the dot product of a predefined template waveform *n* with the raw data is large, while the amplitude of the spike is close to the mean amplitude of template *n*. We thus find local maxima of these dot products over all waveforms and times, and impose a window of ±*t_s_* samples around each of these peaks during which another (smaller) peak cannot be detected, unless the pair of templates corresponding to the peaks have only a negligible overlap (absolute value of dot-product < 0.05). On the final iteration of the spike time inference step, we also perform the template matching step repeatedly, after subtracting off the detected spikes, to find spatiotemporally overlapping spikes (matching pursuit algorithm, see below). To accelerate the learning of the templates, we skip the subtraction on all but the last iteration, and rely on the fact that a majority of spikes from each neuron are identified without resolving spatiotemopral overlaps. Results were very similar when the matching pursuit algorithm was used during the entire optimization procedure.

Finding the spike times, templates and amplitudes is equivalent to minimizing a quadratic function of the form 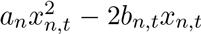 over the scalar variable *x_n,t_*, with *a_n_* and −2*b_n,t_* derived as the coefficients of 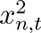 and *x_n,t_* from equation 2:

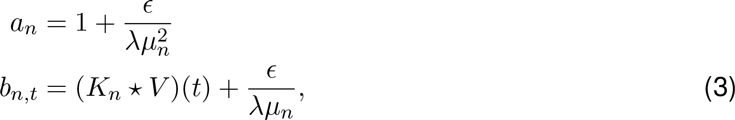

where * represents the operation of temporal filtering (convolution with the time-reversed filter). Here the filtering is understood as channel-wise filtering followed by a summation of all filtered traces, which computes the dot product between the template and the voltage snippet starting at each timepoint *t*. The optimal amplitude *x_n,t_*, and the corresponding decrease in cost *dC*(*n,t*) that would occur if a spike of neuron *n* were added at time *t* are given by:

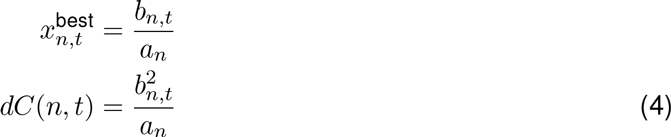

Computing the value of *b_n,t_* for all possible values of *n* and *t* requires filtering the data *V* with all the templates *K_n_*. In principle, this would require a very large number of operations, particularly when the data has many channels. However, the low-rank decomposition of templates allows us to reduce the number of operations by a factor of *N*_chan_/*N*_rank_, where *N*_chan_ is the number of channels (typically > 100) and *N*_rank_ is the rank of the decomposed template (typically 3). This follows from the observation that

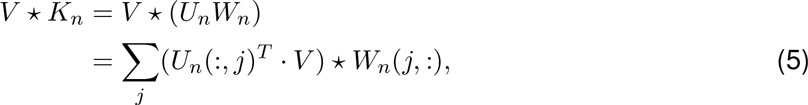

where *U_n_*(:, *j*) is understood as the *j*-th column of matrix *U_n_* and similarly *W_n_*(*j*,:) is the *j*-th row of *W_n_*. We have thus replaced the matrix convolution *V* * *K_n_* with a matrix product 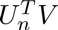 and *N_rank_* one-dimensional convolutions. These matrix products and filtering operations were implemented efficiently using commodity GPU hardware. Iterative updates of *dC* after template subtraction can be obtained quickly using pre-computed cross-template products, as typically done in matching pursuit^25^. The iterative optimization stops when a pre-defined threshold criterion on *dC* is larger than all elements of *dC*.

### *3.3* *Learning the templates via stochastic batch optimization*

The main optimization loop re-estimates the spike times s and template waveforms **K** at each iteration, using a batch-based algorithm to accelerate the optimization and avoid local minima. The data are divided into batches small enough to fit into GPU RAM, and batches are loaded sequentially, in a random order. For each batch, spike times *k* are inferred with the above algorithm, using the current waveform estimates *K_n_*. Then, a running average update rule is run:

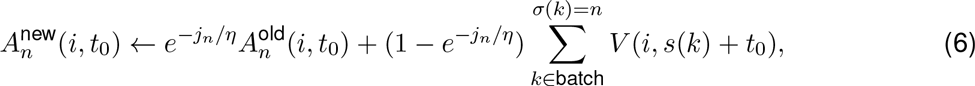

Here, *A_n_* is the running average waveform for cluster *n*, and *j_n_* represents the number of spikes in the current batch identified as belonging to cluster *n* and the running average weighs past samples exponentially with a memory constant *η* (typically ranging from a few dozen to several hundred spikes). Thus, *A_n_* approximately represents the average of the past η samples assigned to cluster *n*. Note that different clusters will update their mean waveforms at different rates, depending on the number of spikes per batch assigned to each cluster. Since firing rates vary over two orders of magnitude in typical recordings (from < 0.5 to 50 spikes/s), this adaptive running average procedure allows clusters with low firing rates to nonetheless average enough of their spikes to generate a smooth running-average template. Periodically during the optimization, we re-estimate the low-dimensional reconstruction of each template from the running average, via an SVD factorization (see section 2.2). The re-estimation is performed every few hundred batches, depending on how large the batches are. In turn, the batch size is limited by the GPU memory and the number of channels sorted together.

Like most clustering algorithms, this model is prone to non-optimal local minima. We used several techniques to ameliorate this problem. First, we annealed several parameters during learning, to encourage exploration of the parameter space, taking advantage of the randomness induced by the stochastic batches. We annealed the forgetting constant *p* from a small value (typically 20 spikes) at the beginning of the optimization to a large value at the end (typically several hundred spikes). We also anneal from small to large the ratio *ϵ*/*λ*, which controls the relative impact of the reconstruction term and amplitude bias term in equation 2; therefore, at the beginning of the optimization, spikes assigned to the same cluster are allowed to have more variable amplitudes. Finally, we anneal the threshold for spike detection (see below), to allow a greater mismatch between spikes and the available templates at the beginning of the optimization. As optimization progresses, the templates become more precise, and spikes increase their projections onto their preferred template, thus allowing higher thresholds to separate them from the noise.

### *3.4* *Determining overlapping spikes in the final inference step*

The main loop alternating template matching and inference is run until the cost function approaches convergence (typically less than six full passes through the data). After convergence, a final inference step is run to detect spatiotemporally overlapping spikes. To find overlapping spikes, we iteratively estimate the best fitting templates (as done in the inference section above), and subtract them off from the raw data. This algorithm is a variant of matching pursuit, which we have parallelized to efficiently implement it on consumer GPU hardware. To see why this parallelization obtains good results, consider the cost improvement matrix *dC*(*n,t*). In standard matching pursuit, when the largest element of this matrix is found and the template subtracted, no values of *dC* need to change except those very close in time to the fitted template (*t_s_* samples away). Thus, instead of finding the global maximum of *dC* like in sequential matching pursuit, we can find local maxima above the threshold criterion, and impose a minimal distance (*t_s_*) between such local maxima. The identified spikes can then be processed in parallel without affecting each other’s representations.

**Figure 3.**
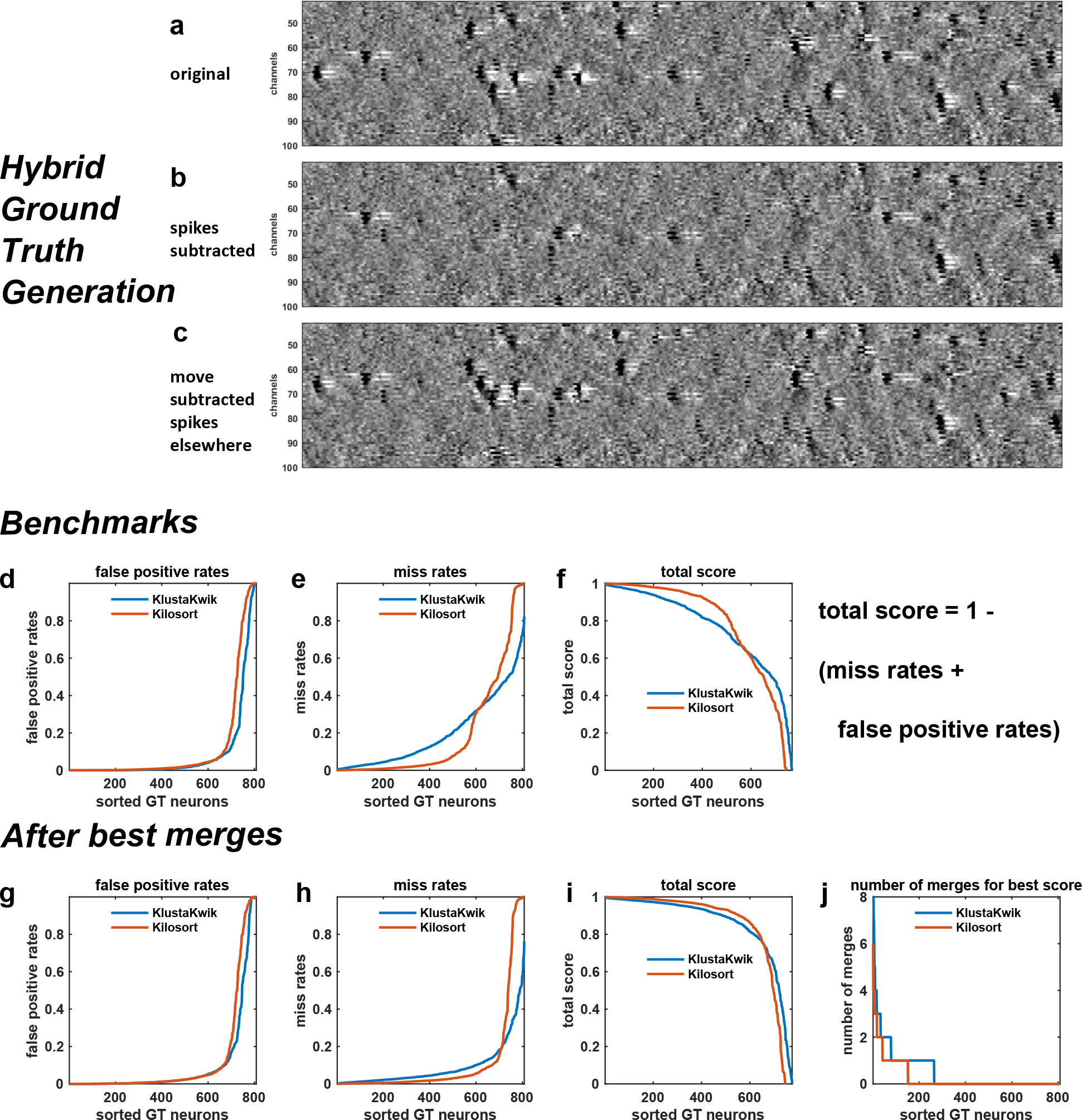
Hybrid ground truth performance of proposed (Kilosort) versus established (KlustaKwik) algorithm. **abc,** Schematic of dataset generation. **d,** False positive rates. **e,** Misses. **f,** Total score. **ghi,** Same as (def) after greedy best possible merges. **j,** Number of merges required to reach best score.

## 4 Benchmarks

To evaluate the algorithm’s performance, we first timed it on several large-scale datasets. The average run times for 32, 128 and 384 channel recordings were 10, 29 and 140 minutes, on a single GPU-equipped workstation (GTX 980Ti). These were significant improvements over an established framework (masked KlustaKwik,^18^), which needed 480 minutes to sort the 32 channel recording, and 15,000 minutes (10 days) to sort the 128 channel recording, while running on a CPU cluster (we did not attempt to run KlustaKwik on 384 channel recordings).

We next asked whether these significant improvements in speed had come at the expense of accuracy. We compared Kilosort and Klustakwik on 32 and 128 channel recordings, using a technique known as “hybrid ground truth”^18^ (Fig. 3a-c). We detail here only the 32 channel benchmarks, which we created by cropping the full 128 channel hybrid benchmark (Fig. 3a-c), which we were able to run with many ground truth neurons. For results on the 128 channel recordings, comparing Kilosort, KlustaKwik and other methods, see http://phy.cortexlab.net/data/sortingComparison/.

To create a hybrid ground truth data set, we first selected all the clusters from a recording that had been previously analyzed with KlustaKwik, and curated by a human expert. For each spike, we extracted its raw waveform and denoised it with an SVD decomposition (keeping the top 7 dimensions, so that waveform variability due to bursting and drift could be modeled accurately). We then added the de-noised waveforms at a different but nearby spatial location on the probe with a constant channel shift, chosen randomly for each neuron. To avoid increasing the spike density at any location on the probe, we also subtracted off the denoised waveform from its original location. This technique therefore produces a dataset with very similar spike waveforms, waveform variability, noise, and synchrony to those found in the original data, but where a large set of spike times are known with 100% accuracy.

Kilosort performed better than KlustaKwik on these hybrids ground truth data, finding more spikes for each of the ground truth cluster (Fig. 3d). We ran both Kilosort and KlustaKwik on 16 instantiations of hybrid ground truth data. To evaluate performance, we matched ground truth neurons with clusters identified by the algorithms to find the maximizer of the score

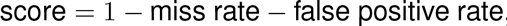

where the false positive rate was normalized by the number of spikes in the test cluster, and the miss rate was normalized by the number of spikes in the ground truth cluster; values close to 1 therefore indicate well-sorted units. Both Kilosort and KlustaKwik performed well (Fig. 3d), with Kilosort producing significantly more neurons with well-isolated clusters (53% vs 35% units with scores above 0.9).

Kilosort also outperformed Klustakwik in terms of best achievable score following manual sorting of the automated results (Fig. 3g-j). Because manually merging an over-split cluster is easier, less time-consuming, and less error-prone than splitting an over-merged cluster, algorithms are typically biased towards producing more clusters than neurons. Both Kilosort and KlustaKwik had such a bias, producing between two and four times more clusters than the expected number of neurons. To estimate the best achievable score after operator merges, we took advantage of the ground truth data, and automatically merged together candidate clusters so as to greedily maximize their score. Final best results as well as the required number of matches are shown in Figure 3g-j (Kilosort vs KlustaKwik 69% vs 60% units with scores above 0.9). The relative performance improvement of Kilosort is clearly driven by fewer misses (Fig 3h), which are likely due to its ability to detect overlapping spikes.

**Figure 4.**
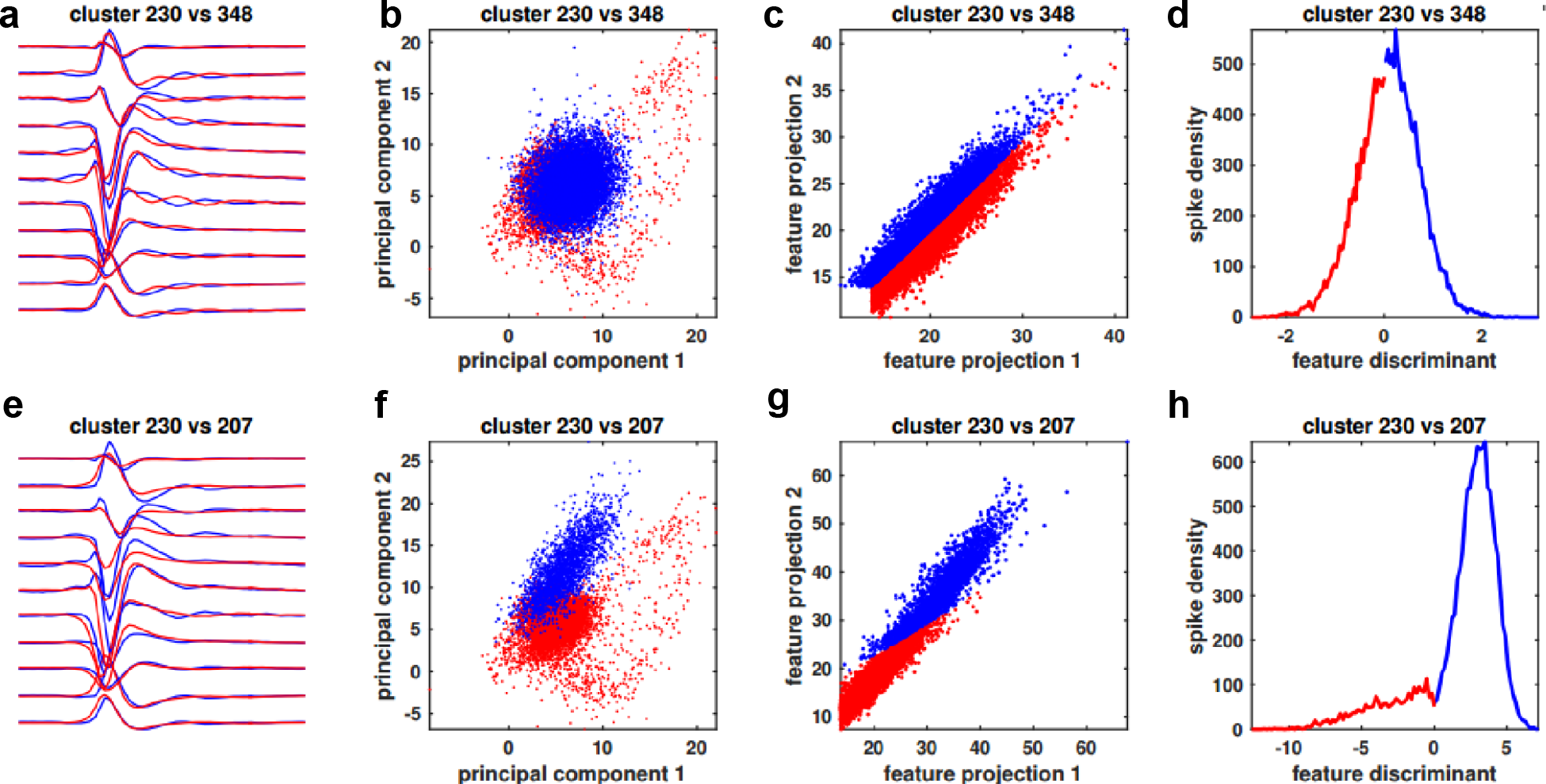
PC and feature-space projections of two pairs of clusters that should be merged. **a,e,** Mean waveforms of merge candidates. **b,f,** Spike projections into the top PCs of each candidate cluster. **c,g,** Template feature projections for the templates corresponding to the candidate clusters. **d,h,** Discriminant of the feature projections from (cg) (see main text for exact formula).

## 5 Extension: post-hoc template merging

We could further reduce human operator work by performing most of the merges in an automated way. The most common oversplit clusters show remarkable continuity of their spike densities (Fig. 4): for such cluster pairs, no discrimination boundary can be identified orthogonal to which the oversplit cluster appears bimodal. Instead, these clusters arise as a consequence of the algorithm partitioning clusters with large variance into multiple templates, so as to better explain their total variance. In Kilosort, we can exploit the fact that the decision boundaries between any two clusters are planes [* To see why the decision boundaries in Kilosort are linear, consider two templates *K_i_* and *K_j_* and consider that we have arrived at the instance of template matching where a spike *k* needs to be assigned to one of these two templates. Their respective cost function improvements are 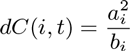, and 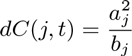, using the convention from equations 3. The decision of assigning spike *k* to one or the other of these templates is then equivalent to determining the sign of *dC*(*i,t*) – *dC*(*j,t*), which is a linear discriminant of the feature projections 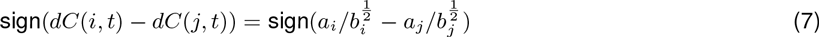 where *b_i_* and *b_j_* do not depend on the data and *a_i, j_* are linear functions of the raw voltage, hence the decision boundary between any two templates is linear (Fig. 4).] to derive an automatic merging heuristic. Indeed, if two clusters belong to the same neuron, their one-dimensional projections in the space orthogonal to the decision boundary will show a continuous distribution (Fig. 4cd and 4gh), and the clusters should be merged. We use this idea to sequentially merge any two clusters with continuous distributions in their 2D feature spaces. Note that projecting both clusters using a single set of principal components for each cluster’s main channel (as would typically be done during manual curation) is much less indicative of a potential merge (Fig 4b and 4f).

### *5.1* *Complete spike-sorting pipeline*

The code is available online at https://github.com/cortex-lab/Kilosort, and can be used together with Phy, a graphical user interface for refining the results of spike sorting (https://github.com/kwikteam/phy; Ref.^18^). We also offer a CPU-based implementation of Kilosort, i. e. one that not requires a GPU, but note that even low-cost commodity GPUs outperform this implementation by at least an order of magnitude.

## 6 Discussion

We have described a new framework for spike sorting of high-channel count electrophysiology data. This framework offers substantial accuracy and speed improvements, while also reducing the amount of manual work required to isolate single units. Kilosort is currently enabling spike sorting of up to 1,000 neurons recorded simultaneously in awake animals (see http://data.cortexlab.net/dualPhase3) and will help to enable the next generation of large-scale neuroscience.

The time taken to run Kilosort scales linearly with the number of recorded neurons, rather than the number of channels, due to the low-dimensional parametrization of template waveforms. Importantly therefore, no performance penalty is incurred by using electrodes of arbitrarily high density.

Although we have defined a heuristic which eliminates the need for many manual merges, operator curation is still required, primarily due to non-stationarities in the recordings such as electrode drift. We anticipate that several strategies combined will reduce this problem. One strategy could involve modeling more explicitly the variability of the templates as a function of time. A second strategy, particularly appropriate for high-density probes, will be to detect the drifts and spatially shift the raw recordings by the inverse of the drifts. Although the drifts cannot be detected from the uniform background activity or the LFP, they can be detected when enough spikes appear to be shifted in one direction more than expected by chance. With such an approach, the possibility of fully automatic spike sorting might soon become reality.

## Acknowledgements

This work was supported by the Wellcome Trust (95668, 95669, 100154), Simons Foundation (325512), and EPSRC (K015141). NS is supported by postdoctoral fellowships from the Human Frontier Sciences Program and the EU Marie Curie program. MC holds the GlaxoSmithKline / Fight for Sight Chair in Visual Neuroscience.

